# Harmonization of CSF and imaging biomarkers for Alzheimer’s disease biomarkers: need and practical applications for genetics studies and preclinical classification

**DOI:** 10.1101/2023.05.24.542118

**Authors:** Jigyasha Timsina, Muhammad Ali, Anh Do, Lihua Wang, Yun Ju Sung, Carlos Cruchaga

## Abstract

**INTRODUCTION:** In Alzheimer’s disease (AD) research, cerebrospinal fluid (CSF) Amyloid beta (Aβ), Tau and pTau are the most accepted and well validated biomarkers. Several methods and platforms exist to measure those biomarkers which leads to challenges in combining data across studies. Thus, there is a need to identify methods that harmonize and standardize these values.

**METHODS:** We used a Z-score based approach to harmonize CSF and amyloid imaging data from multiple cohorts and compared GWAS result using this method with currently accepted methods. We also used a generalized mixture modelling to calculate the threshold for biomarker-positivity.

**RESULTS:** Z-scores method performed as well as meta-analysis and did not lead to any spurious results. Cutoffs calculated with this approach were found to be very similar to those reported previously.

**DISCUSSION:** This approach can be applied to heterogeneous platforms and provides biomarker cut-offs consistent with the classical approaches without requiring any additional data.

## Introduction

Alzheimer’s disease (AD), a degenerative brain disease, is the most common form of dementia and is the sixth leading cause of death in United States ^1^. As the deaths due to other diseases such as stroke, heart diseases etc. continue to decrease, AD related death increased by 71% between 2000-2013 ^1^. Thus, the need for increased focus on understanding the disease and its pathology has become more evident than ever before.

Research suggests AD related brain pathology begins decades prior to onset of clinical symptoms ^2^. Although the diagnosis of AD is based on identification of amyloid plaque and tau tangle accumulation in brain post-mortem, early pathological changes can be tracked using closely related circulating protein biomarkers such as Amyloid (Aβ) and Tau particularly in cerebrospinal fluid (CSF) as well as radio tracers like ^11^C-labeled Pittsburgh compound B or F-florbetapir targeting plaque accumulation in brain using imaging techniques ^3–5^. The use of these endophenotypes is becoming increasingly popular as early detection of AD is of utmost importance given the fact that all current disease treatment modalities are focused on symptom management or disease progression ^6,7^.

The close proximity of CSF to brain, with unrestricted protein flow between the two, and the comparative ease of access to it through lumbar puncture (LP) has made CSF an ideal choice for AD biomarkers among researchers ^8^. Besides the utilization of these proxy CSF biomarkers in AD diagnosis, they can also be used to monitor biological changes throughout the disease progression. Despite these advantages, CSF collection is more invasive when compared to other tissues such as plasma or blood, which has resulted in lower sample availability. This, in turn, has severely limited the ability of researchers to perform large-scale, statistically powerful analysis which could potentially reveal novel AD related pathways and/or biomarkers.

In addition to measuring these circulating protein levels, functional imaging techniques such as positron emission tomography (PET), to identify amyloid plaque and tau accumulation in brain, are also routinely used as a complimentary practice for AD diagnosis ^5,9^. This imaging-based approach allows the non-invasive detection of amyloid and tau aggregates in the brain, a core neuropathologic feature that characterizes AD. There is strong evidence from neuropathologic studies that the most widely used amyloid (i.e., ^11^C-labeled Pittsburgh compound B, ^18^F-florbetapir, and ^18^F-flutemetamol) and Tau PET tracers (i.e., ^18^F-flortaucipir, ^18^F-MK6240, ^18^F-RO948, and ^18^F-PI2620) bind amyloid and tau aggregates, respectively, formed in AD in the more advanced pathologic state (i.e., Braak stage ≥ IV)^10–13^. However, the use of different tracers also creates a problem of combining this data in single analysis.

Since clinical diagnosis of AD is largely based on subjective measurement of cognitive function in patients, efforts have been made to develop a more objective and consistent scale for interpretation of biomarker findings. As a result of these efforts, Clifford et. al. (2016) proposed ATN classification framework which takes three AD biomarkers into account to categorize disease positivity or negativity ^14^. “A” component of the classification describes the Aβ biomarker measured through CSF Aβ level or amyloid PET, “T” refers to Tau, either CSF Tau or Tau PET, and “N” refers to neurodegeneration ^14^. Based on biomarker specific cutoffs, individuals are labelled as either biomarker positive or negative (A+/A-; T+/T-; N+/N-). These classifications are designed to have a consistent and clear format when interpreting results and communicating among clinicians and researchers rather than providing a diagnostic framework.

To increase the quantity and depth of data availability, collaborative research in multi-centers have emerged as a norm to address issues caused by data fragmentation. However, in the absence of an established standards for sample collection and handling as well as measurement techniques and analytical approaches, lack of homogeneity in data has become a glaring obstacle in any data sharing effort. Even when we consider previously described standardization effort such as the ATN classification, the lack of consensus in universal biomarker cutoffs is a major caveat as biomarker level, and subsequently their cutoffs for dichotomization, can be influenced by the technique of measurement. Thus, the need for standard data harmonization approaches exists, particularly those that are focused on endophenotypes such as CSF biomarkers and amyloid imaging among others. One of the special focuses on these endophenotypes is because of their demonstrated utility in genetic studies ^15–17^. Not only have these been used to identify risk variant and genes in context of AD but also in identification of previously unknown disease mechanism. Cruchaga et al., (2013) utilized CSF Tau and pTau in their genome wide association study (GWAS) to identify independent association between these biomarkers and the APOE region. Similarly, by using CSF sTREM2 as endophenotype, Deming et.al., (2019) were able to demonstrate the role of MS4A gene cluster in AD mechanism ^18^. These endophenotypes have also been used to identify sex specific AD risk variants ^19^.

Current practice in terms of data harmonization is focused on re-running previously generated samples using one platform in order to be able to combine samples from different studies. Even with this approach, differences in sample collection can still lead to a technical variation or variation by factors other than the biological nature of the samples called batch effect, which can render the effort of re-running samples useless. In addition, this approach is not practical because of the burden on financial and other resources. Also, with the collection of new samples, the process will have to be repeated. Another approach that is used in genetic studies is meta-analysis, a statistical approach that combines data from multiple independent research geared towards same underlying hypothesis. Doing so is not only tedious as each dataset has to be analyzed individually before it can be meta-analyzed but it also significantly diminishes the power to detect rare variants. As researchers are realizing the importance of conditional and stratified (by sex or disease status) analyses, instead of one size fits all models, these issues become even more cumbersome as they add several time and resource intensive layers to the process. Another major obstacle that we are seeing in data harmonization efforts is lack of consensus among researchers on one standard harmonization approach. As such, the issues that these approaches were designed to address i.e., ease of data sharing and use with consistent replication of results, is still persistent albeit to a lesser degree.

The goal of this paper is to highlight the application of Z-score based data harmonization approaches to CSF biomarker and amyloid imaging. Such approaches would not only help researchers in future collaboration effort but also would allow the use of heterogenous data retrospectively. Increased availability of data is directly translated into statistically powerful genetic studies and better representation of samples, thus making stratified (such as by sex or ethnicity) analysis possible. Z-score has been used previously in genetic studies with great success but its application in data models outside the genetic study framework needs to be evaluated. In this paper, by comparing z-scores with routinely used methods such as Meta-analysis, ATN classification framework, we present evidence that Z-scores based analysis do not lead to spurious results and this method has a potential to be an alternative to traditional approaches. We also demonstrate a data-driven approach to generate cutoffs for ATN classification. Both methods are easy to follow, implement, and are scalable which makes them ideal to be used in collaborative research.

## Methods

### Cohort and Samples

We obtained CSF biomarker levels for Aβ42, Tau and pTau from 23 different cohorts, including Alzheimer’s Disease Neuroimaging Initiative (ADNI) and Knight Alzheimer Disease Research Center (Knight ADRC) cohort among others, encompassing 16,066 total samples that included longitudinal datapoints. CSF measurements were obtained using different platforms such as Elecsys, Lumipulse, Innotest across the cohorts (Supplementary Table 1). Similarly, amyloid imaging data measured using variety of tracers were obtained for 7557 samples (Supplementary Table 2). These subjects were recruited from ADNI, Knight-ADRC, Dominantly Inherited Alzheimer Network (DIAN), Anti-Amyloid Treatment in Asymptomatic Alzheimer’s Disease (A4), ADNI Department of Defense (ADNIDOD), Australian Imaging, Biomarkers and Lifestyle (AIBL), The Harvard Aging Brain Study (HABS) and University of Pittsburgh (UPitt) cohorts. Demographic details of the CSF samples and the amyloid imaging samples is presented in Supplementary Table 1 and Supplementary Table 2 respectively. A subset of the samples was further used to perform GWAS and GMM based cut-off determination analysis.

### Data Quality Control and Standardization

We tested an approach that allows us to combine values from dissimilar cohorts without introducing any batch effect or spurious results. This method is based on the calculation of Z-scores, also known as normal deviate or a standardized score. Z-score shows how many standard deviations (SD) a biomarker value is away from the mean (M) of the dataset ^20^.

Mathematically Z-scores are calculated as,

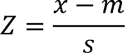

Where, x = observed biomarker value, m = sample mean and s = sample standard deviation. Z-scores can be calculated for each value in a dataset and are easy to interpret as a positive score indicates a raw level above mean and a negative score indicates levels below mean with higher scores showing higher deviation from the average ^21^.

Here, we implemented this approach to 23 cohorts with CSF Aβ42, Tau and pTau level and calculated their Z-scores (Supplementary Table 1). Prior to Z-score calculation, quality control (QC) was done to remove duplicated samples or samples with missing biomarker level. Raw protein values were log transformed followed by outlier removal. Outliers were defined using the Interquartile range (IQR) approach. Any biomarker level lower than Q1-1.5*IQR and higher than Q3+1.5*IQR, where Q1 and Q3 are the first and the third quartile calculated from the distribution, were marked as outlier data points and excluded from the analysis. Outlier detection and removal is a highly recommended step in biomarker-based analysis to exclude technical (e.g., ceiling values) and biological artifacts (e.g., cell lysis) within the dataset to produce robust findings. Z-scores were then calculated using base “scale” function in R statistical software (v3.5.0). The QC steps as well as Z-score calculation were performed for each cohort and each biomarker individually. To demonstrate that the applicability of this framework extends beyond harmonizing CSF specific data types, we employed the same Z-score based harmonization approach for processing amyloid imaging data from the 8 different cohorts. Z-scores were calculated for each cohort and tracer similar to the CSF biomarker datasets as described earlier.

### Comparison against meta-analysis approach

Z-score based data standardization approach has been previously used in several studies focused on identifying genetic variants associated with CSF and amyloid imaging ^15,17,22–25^. However, it is important to demonstrate that this standardization approach does not lead to spurious results. To address this, we performed a joint-GWAS using the CSF Aβ, Tau and pTau Z-scores from 23 cohorts (N =7,231) as phenotype and compared the results with those obtained from meta-analyzing each individual cohort GWAS. First, we used the z-scores obtained from raw biomarker levels from all 23 cohorts, as described earlier, and performed a single variant association analysis for each biomarker, hereby referred to as joint analysis, with these scores as continuous endophenotypes using PLINK v2.0 ^26^. The analysis was adjusted for biological and technical covariates such as age, gender, and genotyping arrays. In parallel and as comparative analysis, each cohort was analyzed independently and the resulting summary statistics were meta analyzed using METAL ^27^, which would constitute the current state-of-the-art approach to combine GWAS result from different cohorts. The inverse variance weighted approach in METAL was used for meta-analysis. Finally, we analyzed the correlation between the effect sizes and p-values from both approaches to determine if both methods would lead to the same results. We implement the same approach for amyloid imaging where we first performed GWAS for eight individual cohorts followed by meta-analysis using METAL and compared the results with those obtained from GWAS using amyloid imaging data z-scores as continuous trait. To further demonstrate the validity of this approach, we asked if conducting an association study using harmonized vs raw endophenotype or log normalized biomarker levels would lead to similar results. We conducted two independent linear regression analyses using amyloid-PET endophenotype as a harmonized (Z-score) and raw Centiloid (CL) scale, available from the Knight ADRC (N = 549) and ADNI (N = 1,134) cohorts and compared their results to check the concordance or lack thereof. We also compared the effect size and p-values from the joint analysis of the eight cohorts using z-scores from amyloid imaging data as phenotype with the meta-analysis result of individual cohort GWAS summary statistics using log transformed biomarker values as phenotypes.

### Unbiased biomarker dichotomization and ATN classification

We used a gaussian mixture model (GMM) to identify z-score driven biomarker cut-off for AT classification of CSF Aβ and pTau from ADNI and Knight ADRC cohorts that used different platform to measure the protein levels. GMM is a probabilistic model approach based on the assumption that all data points within a population can be grouped under a finite number of gaussian distribution thereby identifying subpopulation within them. GMM was implemented through “mclust” package (V 6.0.0) in R statistical software. Within ADNI cohort, xMAP platform was utilized for CSF Aβ measurement and Elecsys platform was used for CSF pTau biomarker measurement. We also leveraged Innotest and Lumipulse data from the Knight ADRC cohort for the measurement of CSF Aβ and pTau levels, respectively. Z-score values for dichotomization were calculated using the same approach as described earlier. The platform-specific cutoffs were determined for each biomarker of interest and each cohort individually, as explained above. From the Z-score cutoff thus identified, the corresponding raw value cutoffs were inferred thereby providing a biologically meaningful biomarker level. We applied the same approach for dichotomizing the amyloid imaging data using different amyloid imaging tracers from ADNI and Knight ADRC. Finally, we assessed the performance of this method in comparison to more classical approaches of biomarker cut-off determination by comparing the agreement of biomarker status determined by our approach within ADNI cohort to that assigned by using previously reported cut-offs for CSF biomarkers. In addition, we also compared the performance of our proposed method with other statistical approach employed for biomarker positivity determination by comparing the agreement between A/T label assigned by each approach in an external dataset.

## Results

### Z-scores transforms dissimilar biomarker levels to a uniform scale

We utilized Z-scores to harmonize CSF Aβ, Tau and pTau biomarker levels from 23 different cohorts measured using various platforms (Supplementary Table 1). The distribution plot of absolute raw biomarkers level and their corresponding Z-scores for Alzheimer’s Disease Neuroimaging Initiative (ADNI) and Knight Alzheimer Disease Research Center (Knight ADRC) shows that the dissimilar raw values are transformed into a uniform Z-score based scale ranging from −3 to 3 in both cohorts (Figure 1A). This is because all biomarker values are standardized with mean 0 and variance 1 regardless of the difference in range of absolute values. The ±3 limits are results of the QC steps we implemented to remove any outlier values prior to Z-score calculation. Similar harmonization and resulting plots were generated for the remaining 21 cohorts as well (Supplementary Figure 1-3). This consensus in scales resulting from z-score standardization allows the use of the biomarkers data from dissimilar sources as one continuous measure. We can also appreciate the fact that the bimodal nature of raw Aβ levels within ADNI cohort, which is an expected observation given the difference between Aβ levels in cases and controls, has been preserved when using Z-score thereby highlighting the ability of Z-scores in capturing underlying biological patterns.

**Figure 1:**
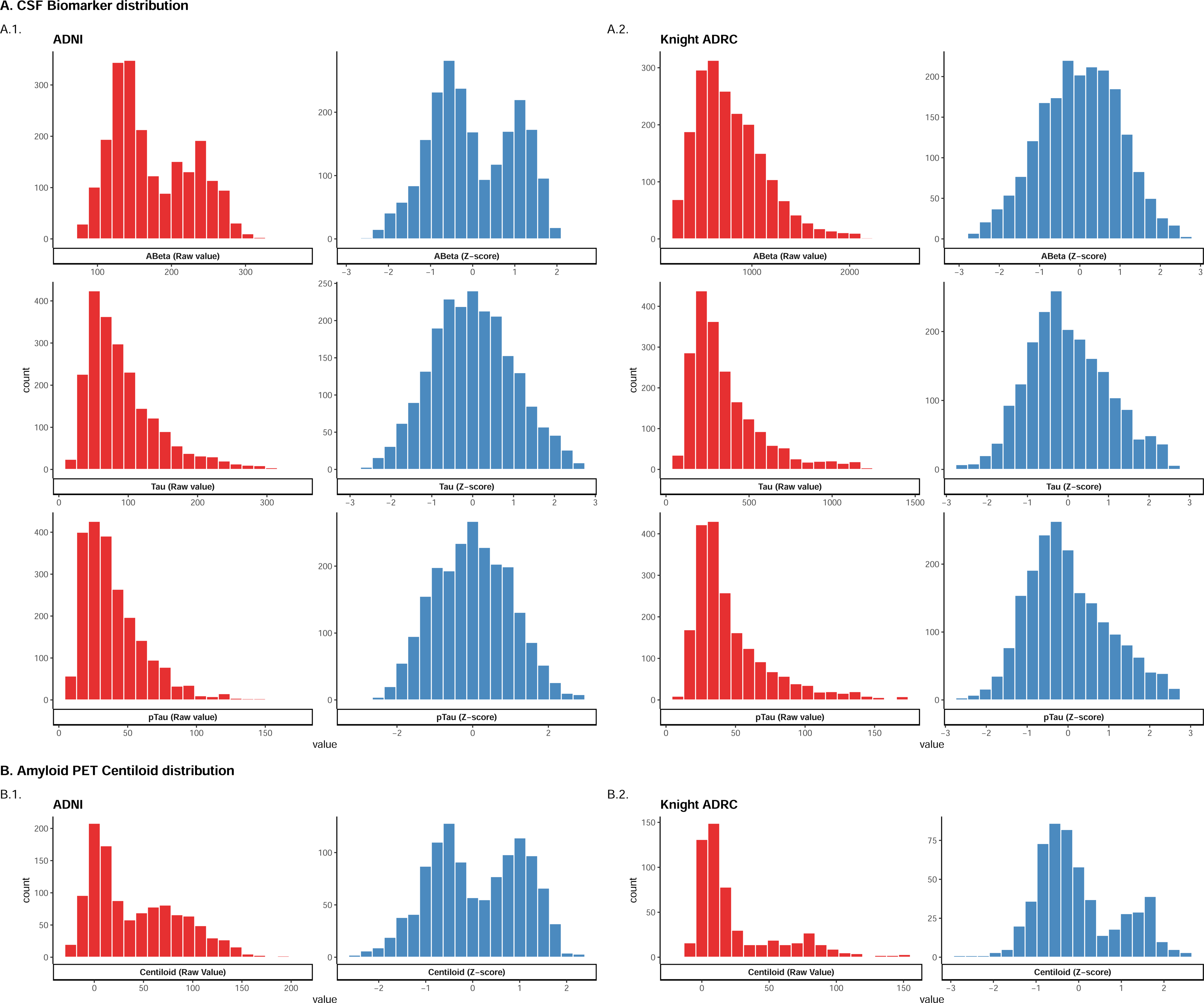
Histogram showing the distribution of raw values and their corresponding Z-score for Aβ, Tau and pTau. (A) and Amyloid PET centiloid values (B) in ADNI and Knight ADRC cohort. Red bins represent raw values and blue bins represent Z-scores The raw values have been assigned a Z-score based on the mean and SD from their distribution in each cohort and scaled to range between −3 to 3. The skewed nature of the raw values was addressed using an intermediate step involving log transformation prior to z-score calculation. Alzheimer’s Disease Neuroimaging Initiative (ADNI); Knight Alzheimer Disease Research Center (Knight ADRC)

To demonstrate the utility of this approach in more than one class of endophenotype, we applied Z-score harmonization approach to amyloid imaging data obtained for 7,557 individuals from 8 different cohorts measured using various tracers (Supplementary Table 2). The standardization approach appropriately overlaid all data sets onto a uniform scale with a mean of 0 and SD of ±3, as seen in the case of CSF biomarker data presented above, regardless of underlying difference in the tracers. Notably, the Z-score transformation preserved the underlying bimodality of the raw amyloid PET data (Figure 1B), suggesting it to be equally favorable for further dichotomizing the harmonized quantitative endophenotype into amyloid-positive and negative populations. Altogether, these observations provide support to the potential utility of Z-scores as a powerful tool in terms of retrospective data harmonization needs.

### Z-score based analysis yields results comparable to meta-analysis

To assess that the Z-score based data standardization does not lead to spurious results, we performed GWAS analysis with z-scores derived using CSF biomarker data (N =7,231) from diverse platform and cohorts, as a continuous quantitative trait and compared the findings with metanalysis of individual cohort GWAS. Both analyses were adjusted for appropriate covariates (Supplementary Figure 4). We found significantly strong correlation between both the effect sizes (r_Aβ_ = 0.969, p < 1×10^-300^; r_tau_ = 0.966, p < 1×10^-300^; r_ptau_ = 0.958, p < 1×10^-300^, Figure 2B) and p-values between the two methods (r_Aβ_ = 0.957, p < 1×10^-300^; r_tau_ = 0.938, p < 1×10^-300^; r_ptau_ = 0.925, p < 1×10^-300^,Supplementary Figure 5) thereby highlighting the utility of z-score based joint analysis as an alternative to meta-analysis.

**Figure 2:**
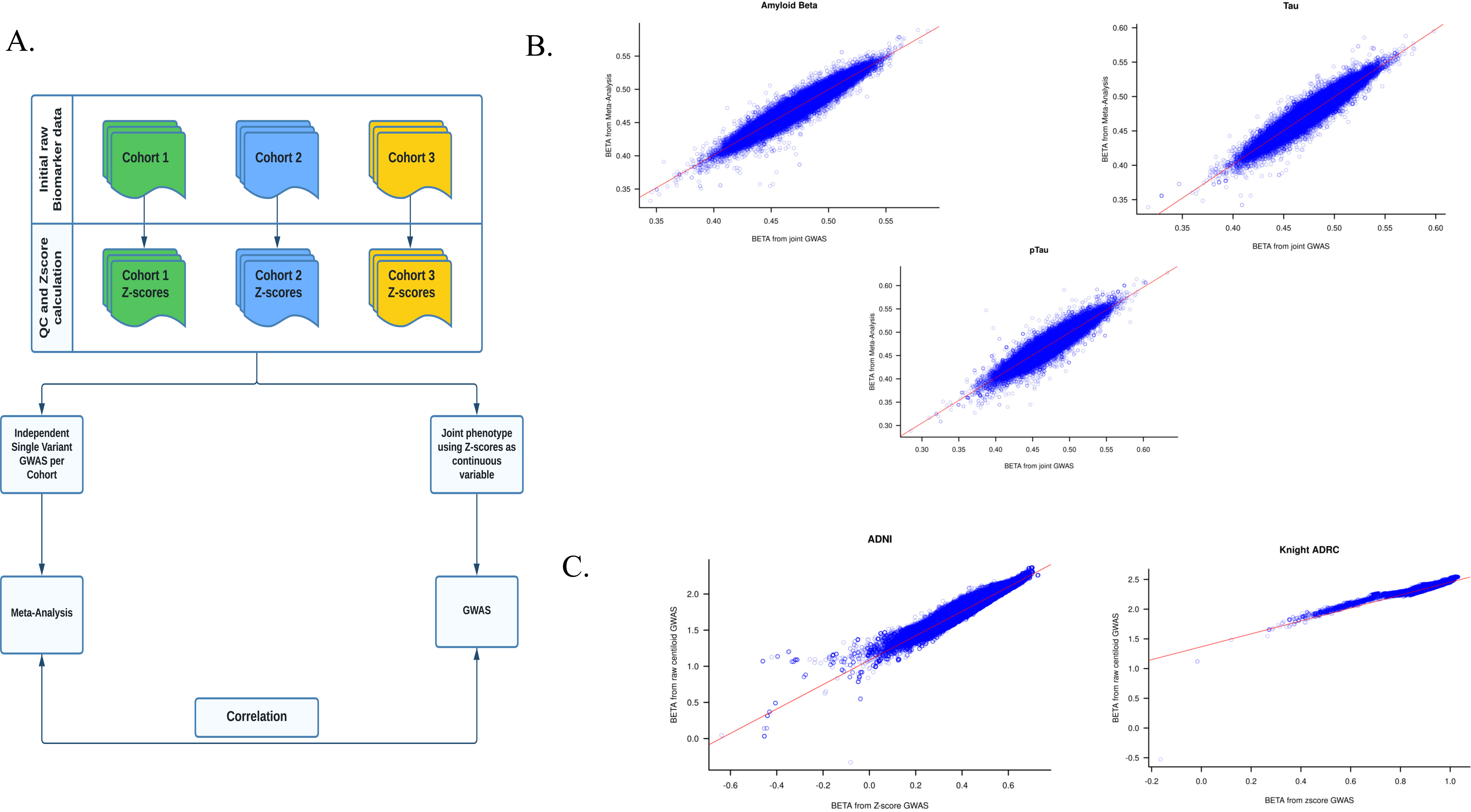
Schematic presentation of Z-score based GWAS result validation process and results. (A)Workflow for joint and meta-analysis GWAS. (B)Correlation between the log10 (effect size) from Z-score based joint GWAS and meta-analysis of individual cohort GWAS for CSF AL (r = 0.969, p < 1×10-300), Tau (r= 0.966, p < 1×10-300) and pTau (r=0.958, p < 1×10-300) (C) Correlation between the log10 (effect size) from Z-score based and raw centiloid values based GWAS for ADNI (r = 0.966, p < 1×10-300) and Knight ADRC (r=0.951, p < 1×10-300).

To demonstrate the utility of this approach in diverse endophenotypes, we applied this harmonization technique to amyloid imaging data obtained for 7,557 individuals from eight different cohorts (Supplementary Table 2) using different imaging tracers (e.g., AV45, FBP, and PiB). As expected, we observed a very high correlation between the p-values (r = 0.980; p < 1×10^-300^; Supplementary Figure 6A-6B) from the joint using amyloid imaging Z-scores as continuous quantitative trait and meta-analysis of individual cohort summary statistics. The strong agreement between joint and meta-analysis approaches highlights the ability of Z-score in data harmonization across different cohorts without producing any false-positive results and increasing the statistical power of the study.

Further, we compared the results from these harmonized values-based analysis with both raw amyloid imaging (Centiloid) endophenotypes based GWAS in Knight ADRC (N = 549) and ADNI (N = 1,134) cohort as well as from meta-analysis of all eight cohorts with log10 transformed biomarker levels as phenotypes. In our analysis comparing results from GWAS with harmonized scores and the raw phenotype, we observed a very strong positive correlation between the effect sizes (r_Knight ADRC_ = 0.951, p_Knight ADRC_ < 1×10^-300^; r_ADNI_ = 0.966, p_ADNI_< 1×10^-300^) and their corresponding p-values (r_Knight ADRC_ = 0.889, p_Knight ADRC_< 1×10^-300^; r_ADNI_ = 0.923, p_ADNI_ < 1×10^-300^) in both the Knight ADRC cohort (Figure 2C; Supplementary Figure 7A) and ADNI cohort (Figure 2C, Supplementary Figure 7B). We found similar strong correlation between results from the joint Z-scores based GWAS and meta-analysis of individual summary statistics using log transformed imaging endophenotype as variable of interest (Supplementary Figure 8). The overall correlation between effect size and p-value for this analysis were 0.98 (p < 1×10^-300^, Supplementary Figure 8) and 0.94 (p < 1×10^-300^, Supplementary Figure 8) respectively.

To summarize, we observed a very strong and significant correlation between joint analysis using Z-scores as phenotype and meta-analysis of individual cohort summary statistics in both CSF and amyloid imaging biomarker. Similar strong correlation was observed when the results from the Z-scores based analysis were compared to results from raw and log transformed amyloid imaging data. These results suggest that the transformation of raw values into a uniform Z-score based scale does not alter the inherent properties of the raw data, thereby, making it an ideal framework for within- and across-cohort data harmonization. Our results also show that using Z-scores as phenotype produces comparable results to log transforming raw endophenotypes. However, since Z-scores based analysis can be performed as single stage analysis by the virtue of uniform variance as a result of the standardization, compared to the two-stage analysis required if using only log transformed values, they become ideal choice when diverse, large-scale data needs to be analyzed.

### Z-score based dichotomization identifies robust biomarker positivity cut-offs

Next, we utilized a GMM based data driven approach using Z-scores to determine biomarker positivity cut-offs without the need of additional information such as amyloid imaging status information and compared the calculated cut-off values with that reported previously for those specific platforms and/or cohort ^28–33^. The assumed biomarker positivity distribution from GMM model is presented in Figure 3. Cut-offs were calculated from both cross-sectional as well as longitudinal data points based on availability. The data-driven cut-off were in agreement with biomarker positivity thresholds reported in the literature (Table 1). For example, in case of the ADNI xMAP CSF Aβ level (N=1244), the data-driven method led to a cut-off of 196 pg/ml, which closely matches with the reported cutoff of 192 pg/ml ^28^. When the analysis was extended to include longitudinal data points (N=2,163), we observed that the cutoff remained consistent with 197 pg/ml being detected as the dichotomization value. In case of CSF pTau biomarker from the Elecsys pTau in ADNI (N=745), our method led to a cut-off of 27.8 pg/ml which is even closer to the one previously reported (27 pg/ml) ^29^. The platform (Innotest) used for measuring CSF Aβ was different in the Knight ADRC cohort (N=1,044) but the estimated cutoff (527 pg/ml) was fairly close to the literature-derived threshold (500 pg/ml) for this platform ^30^. Similarly, for pTau we found a cut-off of 58.9 pg/ml when using cross-sectional data points (N= 1,178) and 58.4 pg/ml with longitudinal data points (N=1,961) in comparison to 58.1 pg/ml previously reported for the platform ^31^. We also applied the same approach for dichotomizing the amyloid imaging data from different tracers and cohorts and obtained similar results (Figure 3; Supplementary Figure 9). For example, in case of Knight ADRC cohort, the amyloid imaging data was obtained using two different tracers (PiB [N=332] and Centiloid [N=494]) but harmonized by the same Z-score based approach. As expected, we observed different amyloid positivity cutoff for PiB (0.12) and Centiloid (33.01), that were in close proximity to what has been previously reported for both these tracers (0.18 and 21.6 respectively) ^32,33^. Overall, the Z-score based empirical cutoff values determined using this mixture model approach showed good agreement with previously reported cutoffs, further validating the practicality of the introduced data harmonization technique.

**Figure 3:**
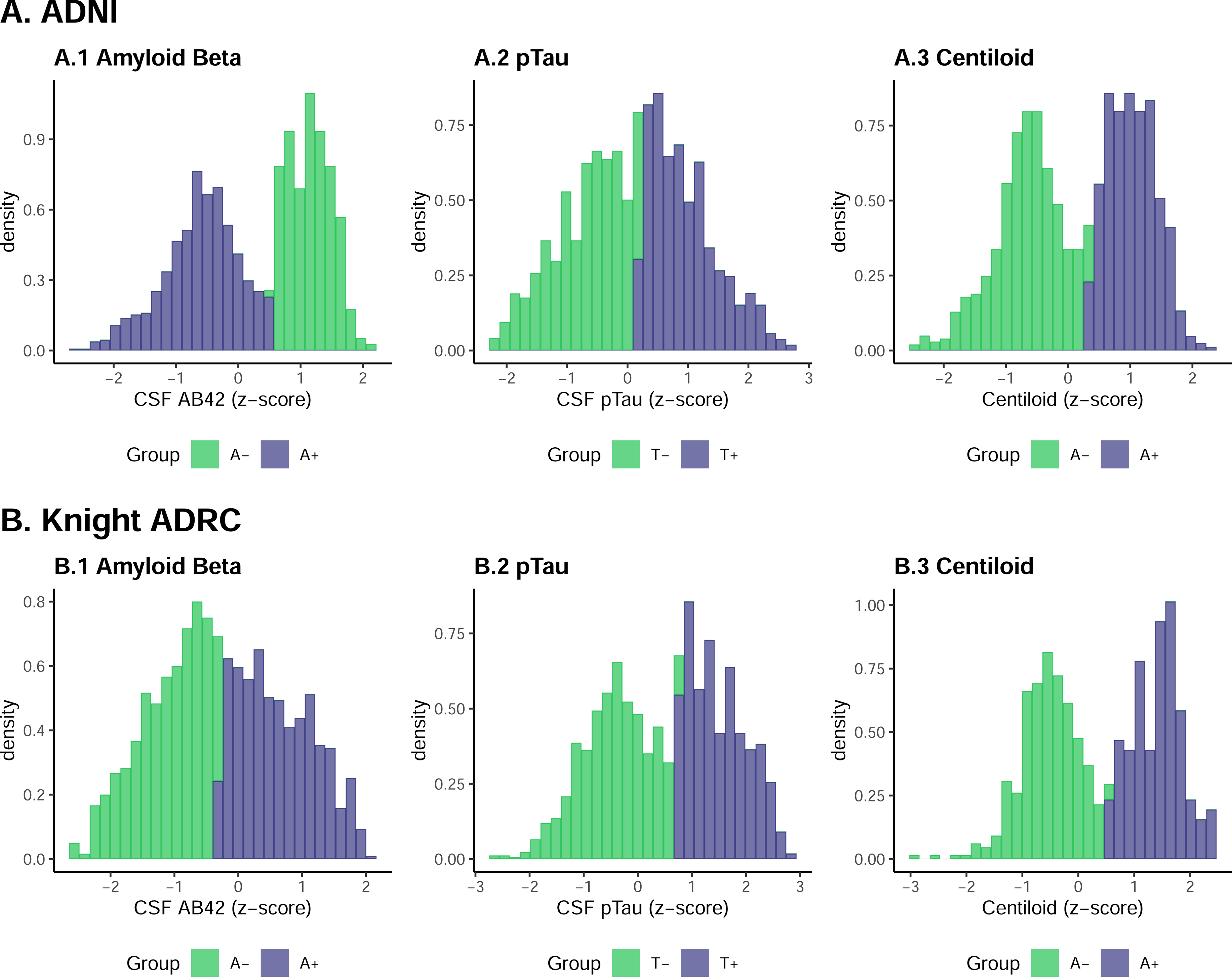
Density plot showing the assumed normal distribution by z-scores based GMM approach in ADNI (A) and Knight ADRC (B) data using CSF biomarkers and Centiloid data from PET imaging. X-axis shows the z-scores values in the dataset. A+/A- and T+/T-denotes biomarker positivity or negativity status. Green bins represent samples assigned biomarker negative (Amyloid/pTau negative) status and bins colored purple represents samples assigned biomarker positive (Amyloid /pTau positive) status.

**Table 1:** The Z-score cutoff and their corresponding raw values determined using Gaussian Mixture Model approach.

| Cohort | Modality | Biomarker | Platform/Tracer | Z-score threshold | Raw value threshold | Reported Raw value threshold |
| --- | --- | --- | --- | --- | --- | --- |
| **ADNI** | CSF (pg/ml) | **Aβ**<br>**(Cross-sectional)** | xMAP | 0.60 | 196 | 192<sup>28</sup> |
|  |  | **Aβ**<br>**(Longitudinal)** | xMAP | 0.57 | 197 | 192<sup>28</sup> |
|  |  | **pTau** | Elecsys | 0.197 | 27.8 | 27<sup>29</sup> |
|  | PET imaging | **Aβ** | Centiloid | 0.43 | 48.93 | 20<sup>33</sup> |
| **Knight ADRC** | CSF (pg/ml) | **Aβ** | Innotest | -0.33 | 527 | 500<sup>30</sup> |
|  |  | **pTau**<br>**(Cross-sectional)** | Lumipulse | 0.73 | 58.9 | 58.1<sup>31</sup> |
|  |  | **pTau**<br>**(Longitudinal)** | Lumipulse | 0.70 | 58.4 | 58.1<sup>31</sup> |
|  | PET imaging | **Aβ** | PiB | -0.29 | 0.12 | 0.18<sup>32</sup> |
|  |  | **Aβ** | Centiloid | 0.55 | 33.01 | 21.6<sup>31</sup> |

Next, we wanted to determine if this method would also lead to comparable ATN classification as reported by other approaches. To this end, we first compared the consistency of biomarker status identified by the mixture modeling in ADNI cohort with those obtained by applying previously reported cutoffs for the biomarkers for the cohort (Table 2). Of the 745 samples with both Aβ42 and pTau levels, 721 samples were assigned same biomarker status by both approaches resulting in an agreement of 96.78% (Table 2). This high concordance between our approach and classical cut-offs provides evidence that the data driven approach can detect reasonable cut-offs for biomarker dichotomization. To further evaluate the performance of our approach, we then utilized CSF biomarker data with known ATN classification of 629 samples, and performed GMM based AT classification for the samples using Z-scores. The previous labels were assigned based on dichotomization cut-offs determined through a Youden’s J index maximization approach ^34^. Our approach determined the Aβ and pTau raw value cutoffs to be 856 pg/ml and 67 pg/ml respectively for this dataset. Two groups with assumed normal distribution were identified by the mixture model using Aβ and pTau individually (Supplementary Figure 10). Using these cutoffs, 219 samples were classified as A- and 412 were labelled as A+. Using pTau cutoffs, 303 samples were assigned to T-status and the rest 327 were assigned to T+ status. When compared to the previously assigned ATN class for these samples, we found a moderate agreement between both approaches with 86.49% of the total samples being assigned same label by both (Table 3). The previously assigned classification did not account for intermediate category like A+/T- or A-/T+. Only 52 samples that were previously assigned as A-were assigned A+ by our approach and 33 samples assigned as T+ previously were assigned as T-. More importantly, no sample classified as A+/T+ were classified as A-/T-, or vice versa. Overall, the concordance observed in both analyses further provides evidence in support of using data driven approaches for dichotomization like the one we have presented here.

**Table 2:** Table showing the comparison between AT classification using GMM and using previously reported cut-offs in ADNI cohort.

|  |  | Reported Cut-offs assigned AT class |  |  |  |
| --- | --- | --- | --- | --- | --- |
|  |  | A-T- | A-T+ | A+T- | A+T+ |
| GMM assigned AT class | A-T- | 224 | 2 | 0 | 0 |
|  | A-T+ | 0 | 25 | 0 | 0 |
|  | A+T- | 5 | 0 | 188 | 16 |
|  | A+T+ | 0 | 1 | 0 | 284 |

**N= 745; "A" represents Aβ status; "T" represents pTau status; "+" symbol denotes a biomarker positivity status; "-" denotes biomarker negative status.**

**Table 3:** Table showing the comparison between AT classification using GMM and previously determined classification in an external validation cohort.

|  |  | Previously assigned AT class |  |
| --- | --- | --- | --- |
|  |  | A-T- | A+T+ |
| GMM assigned AT class | A-T- | 218 | 0 |
|  | A+T- | 52 | 33 |
|  | A+T+ | 0 | 326 |
| N= 629; "A" represents Aβ status; "T" represents pTau status; "+" symbol denotes a biomarker positivity status; "-" denotes biomarker negative status. | | | |

## Discussion

CSF Aβ, Tau and pTau as well as amyloid imaging are among the most established biomarkers in AD research ^11,35^. These have been used in many studies, and are the basis for the current ATN framework which defines the individual’s disease status based on the objective biomarker levels rather than subjective clinical diagnosis scale ^14^. The field of biomarker discovery and study is experiencing exponential growth with more and more proteins being proposed as biomarkers. CSF or plasma neurogranin has been proposed as a cognitive biomarker in CSF and blood exosomes for AD ^36^. NFL reflects neuronal death and is one of the promising blood biomarkers for AD and neurodegeneration in general ^37^. CSF and plasma GFAP, a biomarker for astroglial pathology in neurological diseases ^38,39^ and CSF TREM2 as a biomarker for microglia activation ^18,40^.

Even though more researchers are opting for CSF and PET imaging-based biomarker study, their potential has been stifled by the low sample availability owing to the invasive nature of the CSF sample collection via lumbar puncture and the high cost of the PET imaging. Availability of multiple platforms for biomarker measurement such as Luminex, Elecsys, Innotest and multitude of radiotracers such as ^11^C-labeled Pittsburgh compound B, ^18^F-florbetapir have resulted in heterogenous data that cannot be reliably combined. Even when the same platform is used, the absolute raw biomarker values may still be significantly different between studies because of the differences in sample collection and handling techniques due to lack of a universally accepted protocol. Although plasma biomarkers are emerging, the issues of data heterogeneity that has plagued the field of CSF biomarker and imaging are bound to be perpetuated further with the availability of platforms like C2N ^41,42^ Simoa ^43^ and MSD ^44,45^ among others. For this reason, it is necessary to identify harmonization protocols that allow researchers to combine data across studies and identify individuals that are biomarker positive in an unbiased and consistent manner.

Here, we presented standardized scores as a potential measure for data harmonization. Z-scores are easy to implement and interpret without involving any taxing statistical methodologies and allows researchers to combine biomarker data from different distribution. This is possible because standardizing the values to the mean removes effects introduced by external sources such as different measurement unit or techniques among others. The possibility of combining heterozygous data from diverse settings is of particular importance in case of CSF biomarker research where limited sample size is a consistent challenge ^46,47^. Z-scores can also be used in case of longitudinal data thereby making them scalable in nature. We then demonstrated that these scores can be applied across multiple cohorts and can be used for joint analyses in genetic studies as an alternative to metanalysis without leading to any spurious results. We utilized data from 23 neurodegenerative disease specific cohorts to highlight that joint analysis is comparable to meta-analysis of individual study results. However, joint analysis of standardized values has an edge over meta analyzing independent results from each cohort due to its several advantage. First, joint analysis provides more statistical power for detection of rare variants which may not be detected in individual cohort GWAS. Next, joint analysis allows us to implement more flexible study designs such as sex or disease status stratified analysis in a streamlined manner. By comparing the GWAS summary statistics of imaging data from ADNI and Knight ADRC cohort, we have also shown that these standardized values lead to the same results as raw endophenotypes based analysis. We have successfully used the same approach in multiple studies before ^15,17,25,48^. Deming et.al., (2017) used z-scores to harmonize phenotype data from nine different centers and were able to replicate expected APOE locus in their AD genome wide study in addition to identifying new loci. The replication of *APOE* region provides evidence to support the assumption that z-scores are able to conserve the biological characteristics of the underlying raw biomarker levels. Ali et al., (2022) replicated previously reported association of variants in Klotho gene region and AD through a z-score based GWAS using PET imaging or CSF biomarker data from 17 different cohorts. Similarly, Ali et al., (2023) used z-scores to harmonize the largest amyloid imaging data to date and identified five novel signals associated with brain amyloidosis. Taken together, the Z-score transformation provides an ideal framework for normalizing the within- and across-cohort variation in the raw endophenotypic data without masking its inherent biological characteristics. One may argue the some of the positive attributes of z-scores highlighted here are shared with the log transformation performed prior to the standardization, but the uniformity of variance across dissimilar dataset that allows for the use of the diverse biomarker data as a continuous trait is inherent to Z-score and cannot be achieved through log normalization in itself.

We have also extended the use of standardized scores to identify the cut-off point that defines biomarker positive from negative using a GMM approach. Here we demonstrate that using this approach leads to very similar cut-off values to those reported using more classical approaches (Table 1). This can be leveraged to perform the ATN classification at individual level, which leads to also very consistent and replicable results, as shown by our analysis comparing the ATN labels assigned by the method with previously identified labels in an external cohort. Current ATN classification approaches are calculated per dataset and are based on comparing a specific biomarker; such as CSF Aβ, with a different standard (for example amyloid imaging) to identify the cut-off for biomarker positivity. This approach requires to have multiple comparable biomarkers for the same individual and in a large number of samples which make these analyses more complicated. Data driven approaches such as the one presented here could be an alternative approach to identify the cut-off point to define biomarker positivity without the need of additional markers. The real value of this approach is in use cases where we do not have any information on the platform used for biomarker measurement, when tested in a new population, when a new assay or platform is being used or to determine gender or race specific cut-off. Several studies have previously reported difference in biomarker levels between these population subgroups ^49,50^, therefore this approach can be easily implemented without the need to have access to additional data.

Despite these demonstrated strengths, there are some limitations to this method. One potential critique of this method is that by using standardized values, the mean of the distribution becomes or approximates 0 which is not informative and does not correspond to any biological value or biomarker cut-off. The mean of any marker distribution is a property of the cohort and is representative of the cohort characteristics. For example, in a cohort enriched for AD cases compared to controls, the average CSF Aβ will shift to the left in comparison to the cut-off for biomarker positive and in a cohort with more controls the mean will be shift to the right. As such, by setting the mean to be 0 we are losing this biological information thereby providing credibility to the argument. However, this can be addressed by combining the standardized values with the dichotomization approach presented here. First, the raw biomarker values and standardized values can be calculated as shown here. This gives a distribution with a mean of 0 and standard deviation of 1. However, in contrast to other methods, such as rank based scores, this method preserves the overall distribution of protein levels. Second the cut off for biomarker positivity is calculated using the same GMM approach as presented here. As shown by our results here, this data-driven method does not require any other information to identify the cut-off and is highly replicable. Third, once the cut-off is calculated, new values can be re-calculated by centering the raw values using the cut-off. In this way, for all the different studies and biomarkers, 0 will be always the point of separation between biomarker positive and negative individuals. Finally, as all datasets will have similar standard deviation and are centered around the cutoff, the value for each individual will be highly informative as it will inform whether the value corresponds to biomarkers positive and negative status, and how far are from the cut off or how extreme it is even though these values do not correspond to any absolute values in protein levels and cannot be represented in pg/ml or any other scale. This has the advantage that there will not be a need to have specific cut off values for each assay. However, since the resulting distribution is not Z-score distribution, the biological feature and utility of the new distribution as a proxy will need further evaluation using similar approaches as presented in this paper for z-score distribution. Another limitation of this method is that in order to identify a reliable cut-off, we need a sample size of at least 200 individuals with a relatively good balance of both cases and controls. However, as more and more biomarker and imaging data becomes available, we foresee that this would not be a limitation in near future.

Currently there is an effort to perform harmonization of multiple endophenotypes (cognition, raw imaging, neuropath, CSF biomarkers) for individuals included in the ADSP study and We harmonization of the CSF biomarkers using the standardized values as well as the ATN classification for all cohorts with available biomarkers data. We are depositing our results in NIAGADS (<u>Alzheimer’s Disease Sequencing Project Phenotype Harmonization Consortium (ADSP-PHC) – ADSP (niagads.org)</u> as part of the U24 harmonization consortium. We are currently implementing the additional harmonization as described here, and expect to release the data in near future.

## Conclusion

In conclusion, to address the need of data harmonization in the field of CSF biomarker and amyloid imaging field, we presented a Z-score-based approach. Not only does this method allows the combination of data from dissimilar platforms and studies, but also preserves the characteristics of underlying raw data and produces comparable results in genetic studies. In addition, when used in combination with mixed modelling approach, they are helpful in identifying biologically relevant cutoffs for distinguishing biomarker positive and negative individuals that are close to reported cutoffs currently being used. Overall, Z-score based method can be a powerful solution for data harmonization.

## Consent Statements

All applicable informed consent were obtained from study participants by individual cohorts as such no new consent was necessary for this project.

