## Supplementary Materials for "Harmonization of CSF and imaging biomarkers for Alzheimer’s disease biomarkers: need and practical applications for genetics studies and preclinical classification"

| Supplementary Table 1: Demographic information of samples with CSF biomarkers information | | | | | | | | | | | |
| --- | --- | --- | --- | --- | --- | --- | --- | --- | --- | --- | --- |
| Cohort | **Platform** | **N** | **Average Age**  **(SD)** | **%Male** | **Average A42** | **Average Tau** | **Average pTau** | **%Cases** | **%Controls** | **%Others** | |
| ADNI | xMAP | 2174 | 74.71 (±7.24) | 56.35 | 172.88 | 92 | 39.93 | 69.04% | 30.40% | | 0.55% |
| AIBL | Innotest | 423 | 73.81 (±6.27) | 52.01 | 826.03 | 302.26 | 58.99 | 20.09% | 46.81% | | 33.10% |
| BIOCARD | NA | 761 | 59.98 (±10.29) | 41.92 | 409.24 | 71.09 | 38.69 | 6.04% | 91.72% | | 2.23% |
| Blennow | NA | 78 | 82.26 (±10.54) | 29.49 | 560.92 | 659.71 | 72.85 | 53.85% | 46.15% | | 0% |
| Barcelona-1 | ELISA | 301 | 67.54 (±8.51) | 49.17 | 931.32 | 460.59 | 69.54 | 23.59% | 1.33% | | 75.08% |
| DIAN | Lumipulse | 1010 | 40.03 (±10.93) | 42.48 | 634.27 | 440.48 | 61.45 | 26.14% | 73.86% | | 0% |
| DOD | Elecsys | 144 | 68.92 (±3.9) | 99.31 | 1244.79 | 216.95 | 19.14 | 27.78% | 68.05% | | 4.17% |
| FACE | NA | 632 | 72.19 (±8.41) | 41.61 | 767.68 | 515.87 | 79.6 | 57.12% | 27.22% | | 15.66% |
| HB | NA | 107 | 67.23 (±9.38) | 53.27 | 75.65 | 85.4 | NA | 100% | 0% | | 0% |
| Lleo | NA | 144 | 64.13 (±9.69) | 34.72 | 625.56 | 413.66 | 59.05 | 36.81% | 50.69% | | 12.50% |
| London | NA | 315 | 68.66 (±9.09) | 53.65 | 390.55 | 100.14 | 43.6 | 61.59% | 9.21% | | 29.20% |
| MAP | Lumipulse | 1999 | 69 (±9.2) | 43.62 | 790.27 | 356.79 | 46.52 | 19.36% | 80.59% | | 0.05% |
| MARS | NA | 103 | 64.69 (±8.67) | 59.22 | 834.88 | 222.38 | 37.96 | 84.47% | 0% | | 15.53% |
| MAYO | NA | 579 | 78.87 (±6.15) | 61.49 | 326.58 | 102.09 | 22.06 | 21.59% | 77.03% | | 1.38% |
| Moli | NA | 256 | 64.61 (±8.12) | 36.72 | 574.03 | 470.89 | 78.59 | 67.19% | 21.09% | | 11.72% |
| NACC | Multiple | 2558 | 71.67 (±9.19) | 49.14 | 469.85 | 308.93 | 51.37 | 33.62% | 24.63% | | 41.75% |
| PPMIS | NA | 2964 | 62.76 (±9.98) | 63.33 | 932.93 | 176.2 | 15.51 | 54.69% | 25.37% | | 19.94% |
| SWEDEN | NA | 315 | 75.13 (±7.6) | 36.83 | 262.59 | 777.34 | 105.38 | 100% | 0% | | 0% |
| UPENN | Luminex | 256 | 71.64 (±8.89) | 43.75 | 167.67 | 94.63 | 37.34 | 64.06% | 18.75% | | 17.19% |
| UW | NA | 441 | 60.22 (±17.35) | 50.57 | 143.23 | 60.5 | 55.58 | 31.52% | 63.26% | | 5.22% |
| VMAP | NA | 156 | 72.4 (±6.33) | 67.31 | 712.63 | 423.74 | 61.43 | 37.18% | 53.21% | | 9.61% |
| WiscADRC | NA | 270 | 63.99 (±10.09) | 38.15 | 669.15 | 384.25 | 50.27 | 13.70% | 75.93% | | 10.37% |
| Zetter | NA | 80 | 73.36 (±4.05) | 50 | 617.15 | 428.86 | 75.94 | 40% | 25% | | 35% |
| NA= Not available; N= Number of samples; Age = Age at CSF draw in years; SD= Standard Deviation. Cases include AD cases only; Others include samples with diagnosis other than AD like FTD, DLB or unknown status. | | | | | | | | | | | |

| Supplementary Table 2: Demographics of amyloid PET GWAS participants at the time of scanning | | | | | | | | | |
| --- | --- | --- | --- | --- | --- | --- | --- | --- | --- |
| **Variables** | **Total** | **A4** | **ADNI** | **AIBL** | **ADNI**-**DOD** | **DIAN** | **HABS** | **Knight-ADRC** | **UPitt** |
| **Tracer** | - | SUVR | FBP | FBP/PIB/ FMT | AV45 | PIB | PIB | PIB/AV45 | PIB |
| **Total** | 7,557 | 3,180 | 1,134 | 1,214 | 169 | 209 | 258 | 1,048 | 345 |
| **Female (%)** | 46.78 | 60.09 | 47 | 54.04 | 0.59 | 52.63 | 59.3 | 54.77 | 45.8 |
| **Male (%)** | 53.22 | 39.91 | 53 | 45.96 | 99.41 | 47.37 | 40.7 | 45.23 | 54.2 |
| **Age (mean)** | 68.31 | 71.30 | 73.73 | 72.65 | 69.08 | 36.60 | 73.99 | 70.74 | 78.39 |
| **Age (sd)** | 7.44 | 4.73 | 7.62 | 6.62 | 4.67 | 10.86 | 6.1 | 9.01 | 9.91 |
| ***APOE*4+ (%)** | 34.15 | 36.04 | 43.39 | 35.01 | 26.63 | 28.71 | 29.84 | 41.13 | 32.46 |
| **Cases (%)** | 25.37 | 0.06 | 61.55 | 7.5 | 32.54 | 45.01 | 27.13 | 16.98 | 12.17 |
| **Controls (%)** | 67.54 | 99.94 | 32.36 | 79.74 | 67.46 | 38.28 | 72.87 | 61.83 | 87.83 |
| For each cohort, we report number of participants, percentage of females and males, mean age of the participants and standard deviation (SD) in the age, percentage of *APOE* ε4-carriers (*APOE* ε4+ participants), and percentage of cases and control participants, where available. To normalize amyloid PET endophenotype across different cohorts, we converted different amyloid imaging measures (e.g., Centiloid, PiB, and AV45) into log-normalized z-score using “scale” function in base R. Phenotype from each cohort was normalized individually to account for within cohort variation. Abbreviations: PET, positron emission tomography; sd, standard deviation; Knight-ADRC, Knight Alzheimer’s Disease Research Center ; ADNI, Alzheimer's Disease Neuroimaging Initiative; DIAN, the Dominantly Inherited Alzheimer Network; A4, Anti-Amyloid Treatment in Asymptomatic Alzheimer's Disease; ADNI-DOD, ADNI Department of Defense studies; AIBL, Australian Imaging, Biomarkers and Lifestyle; HABS, The Harvard Aging Brain Study; UPitt, University of Pittsburgh; SUVR, standardize uptake value ratios; FBP, Florbetapir; PIB, Pittsburgh Compound-B; FMT, Flutemetamol. | | | | | | | | | |

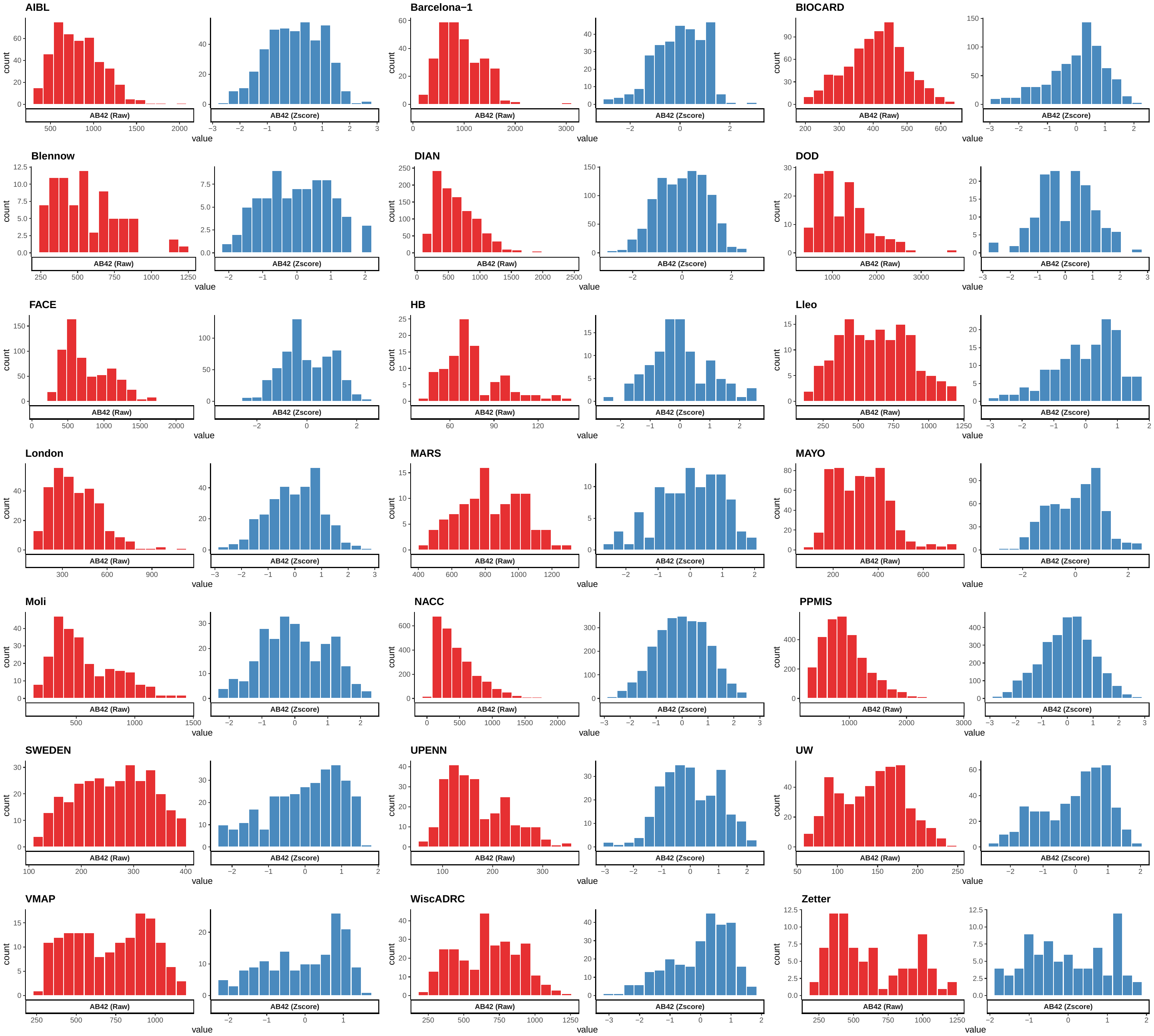

**Supplementary Figure 1: Histogram showing raw and z-score normalized CSF Amyloid Beta levels in individual cohorts**. The raw values have been assigned a Z-score based on the mean and SD from their individual distribution and scaled to range between -3 to 3. Raw and Z-score values are presented in x-axis and data frequency is presented in y-axis. Red bins represent raw values and blue bins represent Z-scores.

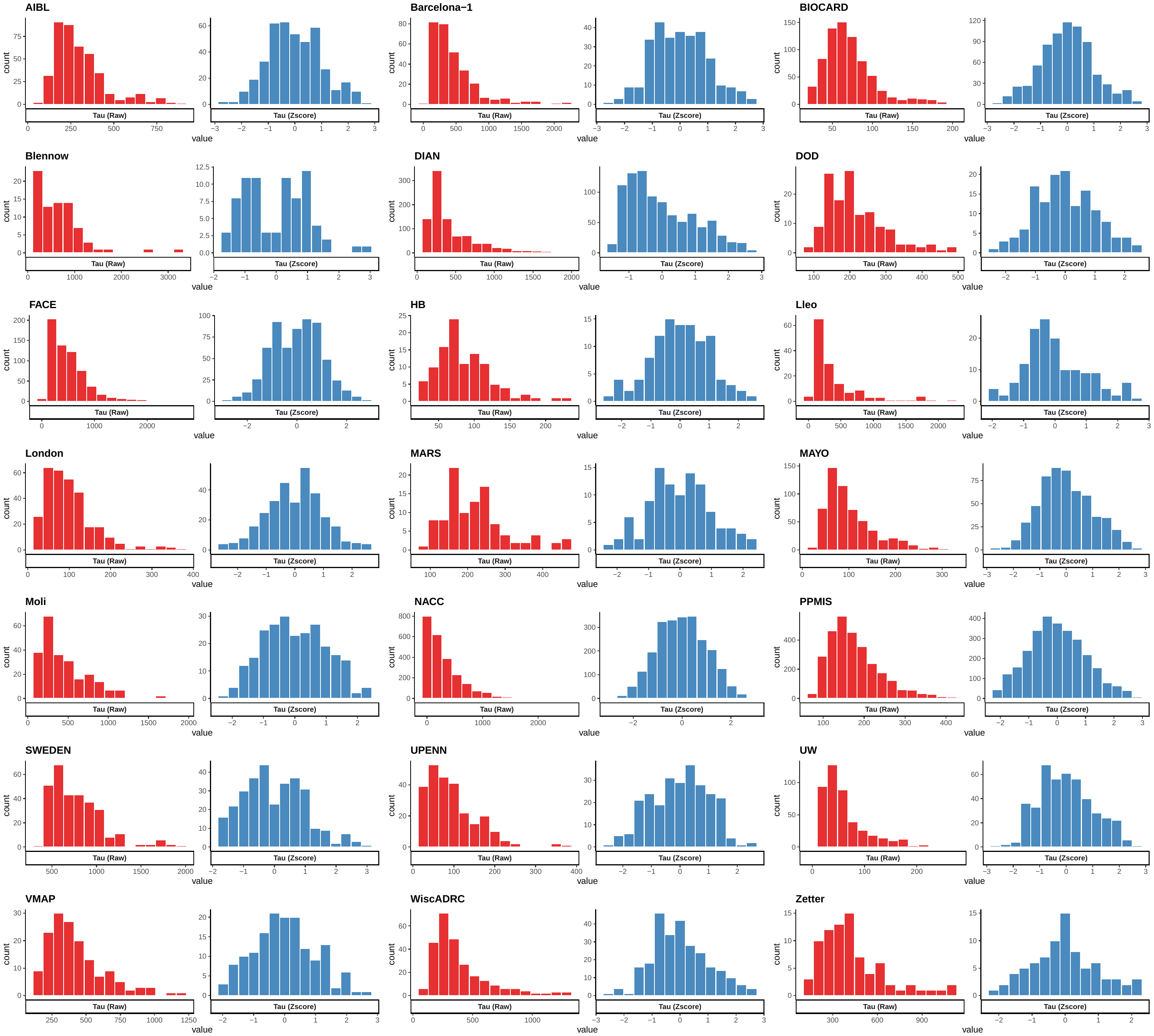

**Supplementary Figure 2: Histogram showing raw and z-score normalized CSF Tau levels in individual cohort**. The raw values have been assigned a Z-score based on the mean and SD from their individual distribution and scaled to range between -3 to 3. Raw and Z-score values are presented in x-axis and data frequency is presented in y-axis. Red bins represent raw values and blue bins represent Z-scores.

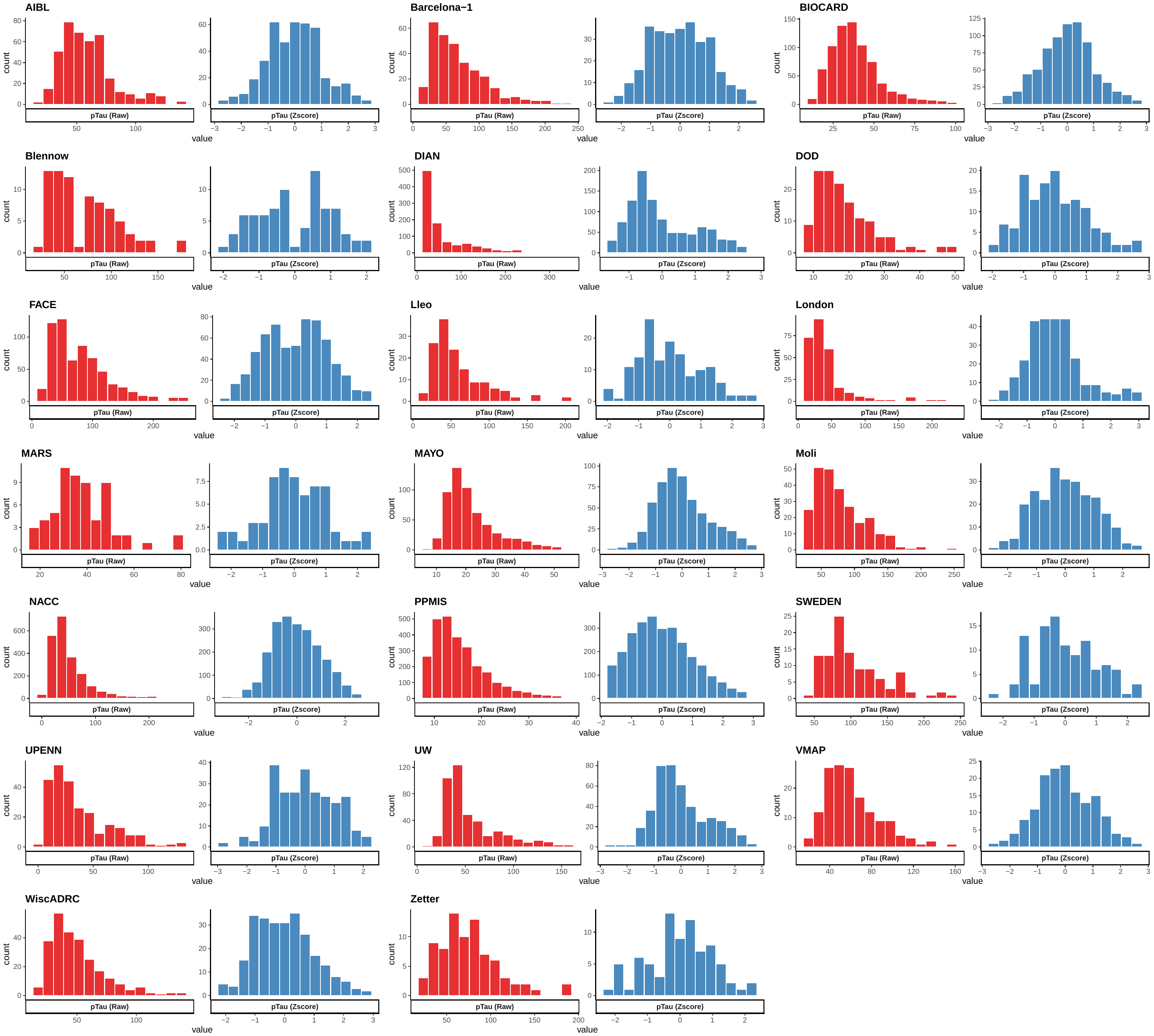

**Supplementary Figure 3: Histogram showing raw and z-score normalized CSF pTau levels in individual cohort**. The raw values have been assigned a Z-score based on the mean and SD from their individual distribution and scaled to range between -3 to 3. Raw and Z-score values are presented in x-axis and data frequency is presented in y-axis. Red bins represent raw values and blue bins represent Z-scores.

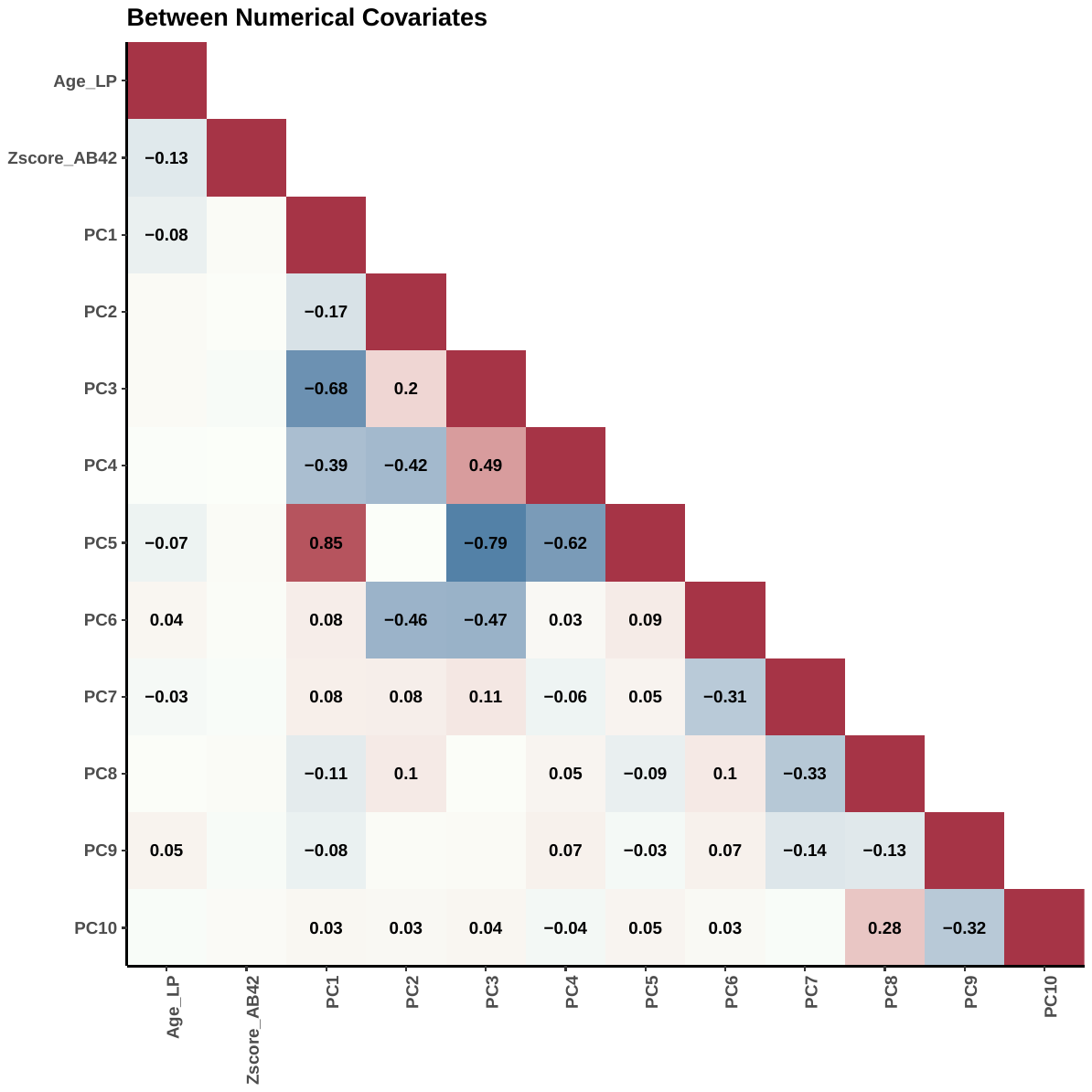

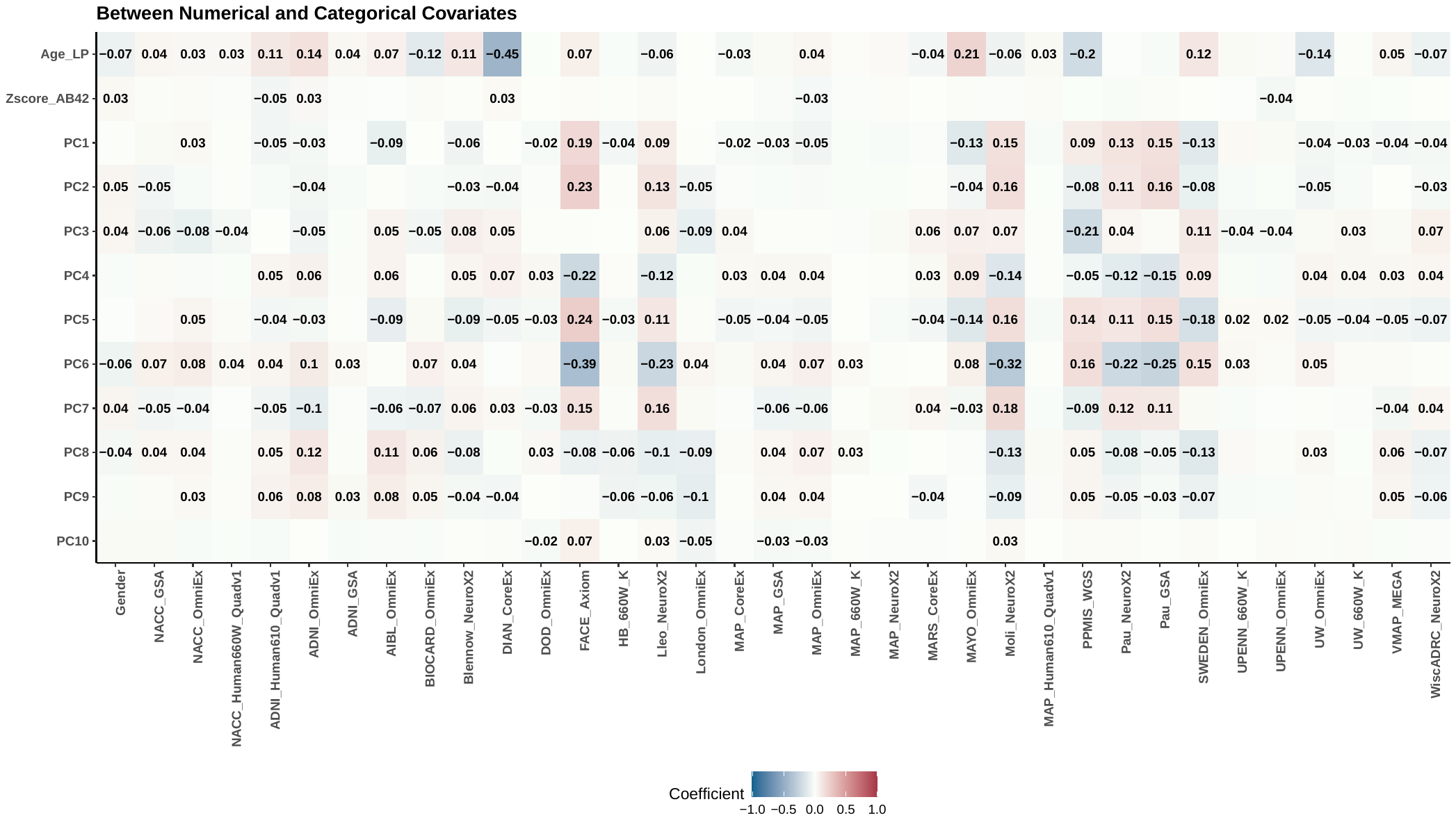

**Supplementary Figure 4:Correlation matrix showing correlation coefficient between one of the CSF biomarkers tested in GWAS analysis and covariates used.** Only those correlations that passed significance threshold (p < 0.05) are shown. Correlation between numerical variable determined using Pearson’s method. Correlation between categorical and numerical variable determined using point-biserial approach. Highest positive correlation was found between PC1 and PC5 with a coefficient of 0.85 and p –value < 10^-300^. The highest negative correlation was found between PC3 and PC5 with correlation coefficient of –0.79 and p –value < 10^-300^.

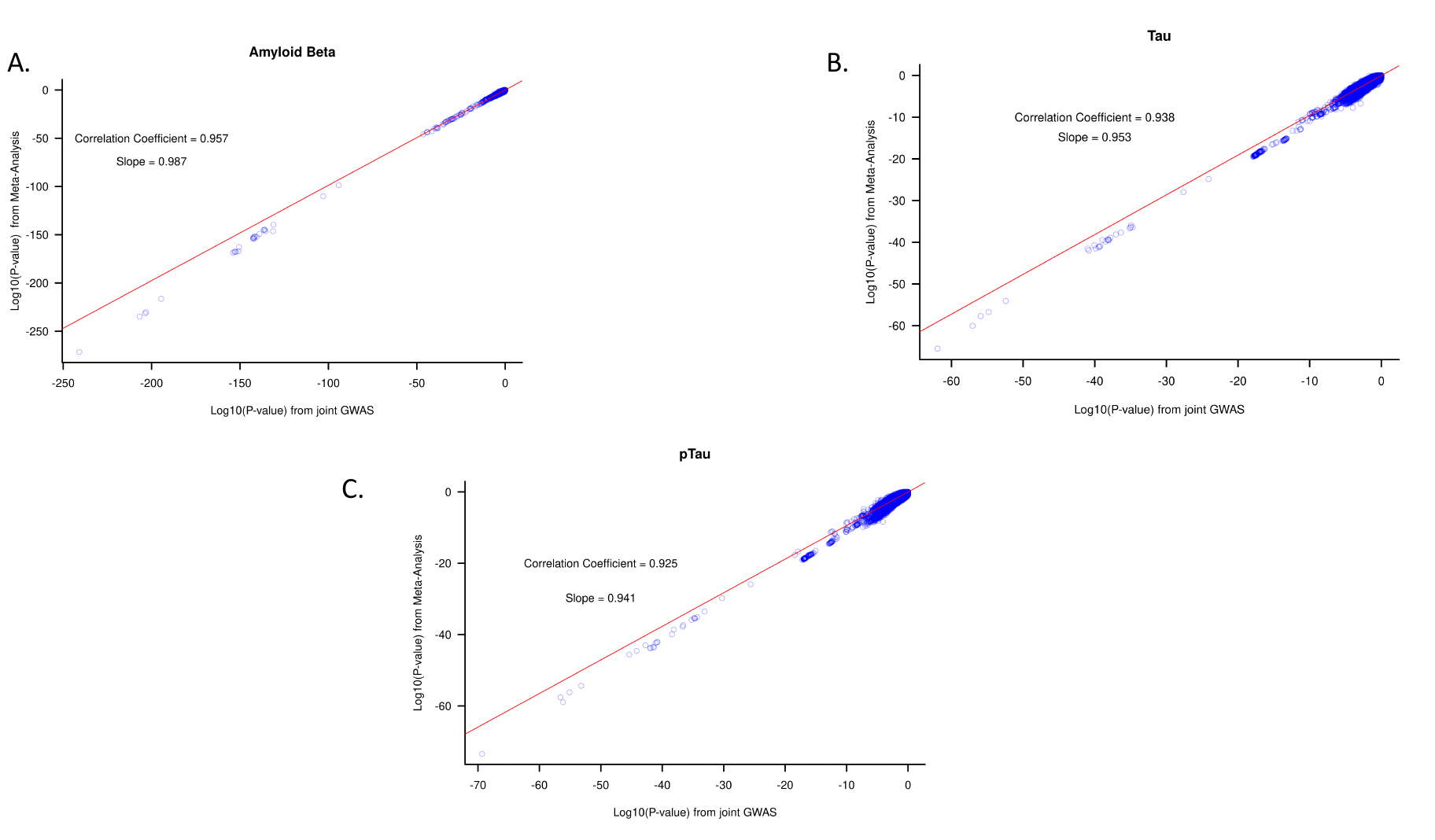

**Supplementary Figure 5: Correlation between P-values from CSF biomarker GWAS from 23 cohorts** A) Plot showing the correlation between P-values from the standard-error (StdErr)-based meta-analysis by METAL software (Y-axis) and Joint analysis (X-axis) from the GWAS analysis using CSF Aβ42 zscores as phenotype. B) Plot showing the correlation between P-values from the standard-error (StdErr)-based meta-analysis by METAL software (Y-axis) and Joint analysis (X-axis) from the GWAS analysis using CSF Tau zscores as phenotype. C) Plot showing the correlation between P-values from the standard-error (StdErr)-based meta-analysis by METAL software (Y-axis) and Joint analysis (X-axis) from the GWAS analysis using CSF pTau zscores as phenotype.

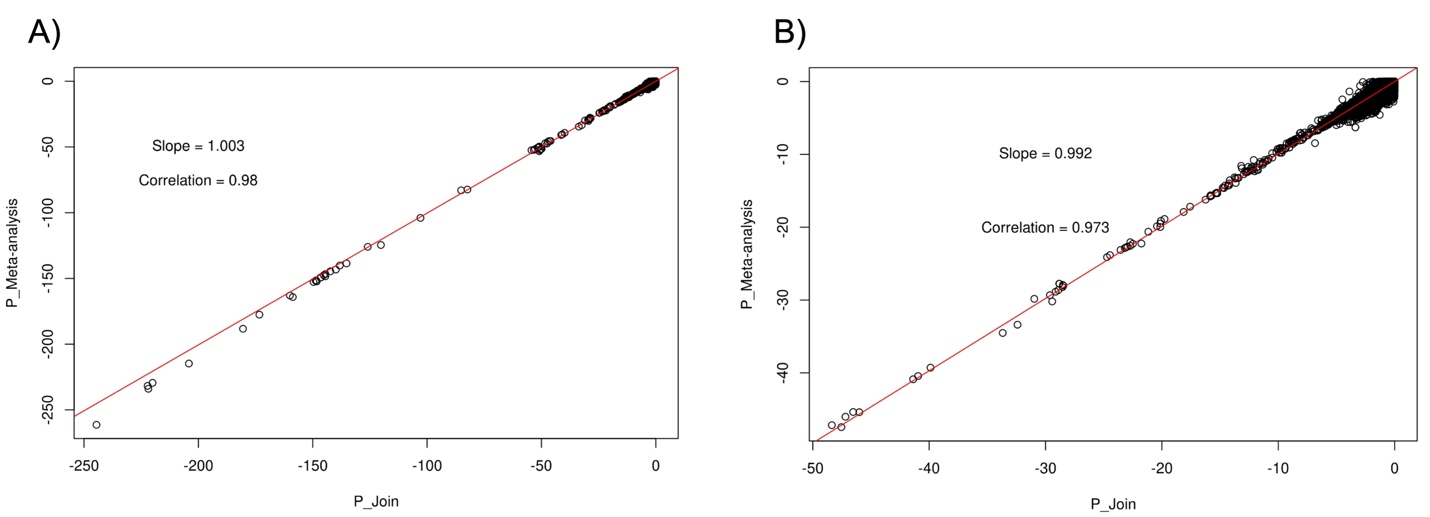

**Supplementary Figure 6: Correlation between P-values from amyloid imaging GWAS from 8 cohorts** A) Plot showing the correlation between P-values from the standard-error (StdErr)-based meta-analysis by METAL software (Y-axis) and Joint analysis (X-axis) from the GWAS analysis using amyloid imaging zscores as phenotype from eight different cohorts. B) Same correlation plot as shown in panel A but the P-values were restricted to P-values greater than 2 × 10−50.

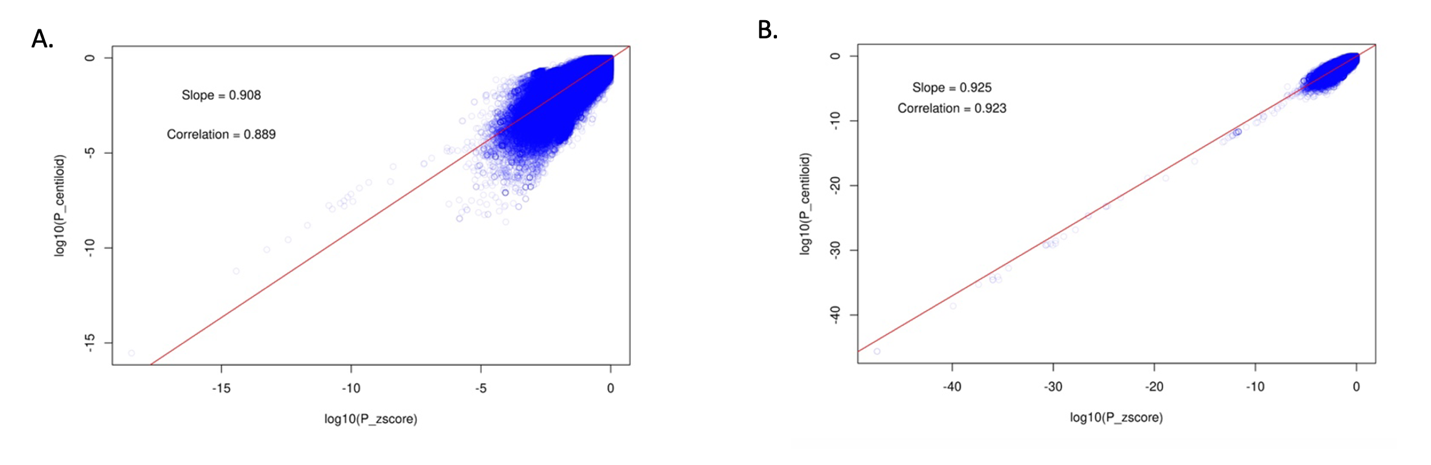

**Supplementary Figure 7: Correlation between P-value for GWAS using Z-score and raw Centiloid measure as the phenotype from Knight-ADRC and ADNI cohorts.** Correlation between the P-value (A) from the GWAS analyses using Z-score (x-axis) and Centiloid measure (y-axis) as phenotype in Knight-ADRC cohort. Correlation between the P-value (B) from the GWAS analyses using Z-score (x-axis) and Centiloid measure (y-axis) as phenotype in ADNI cohort.

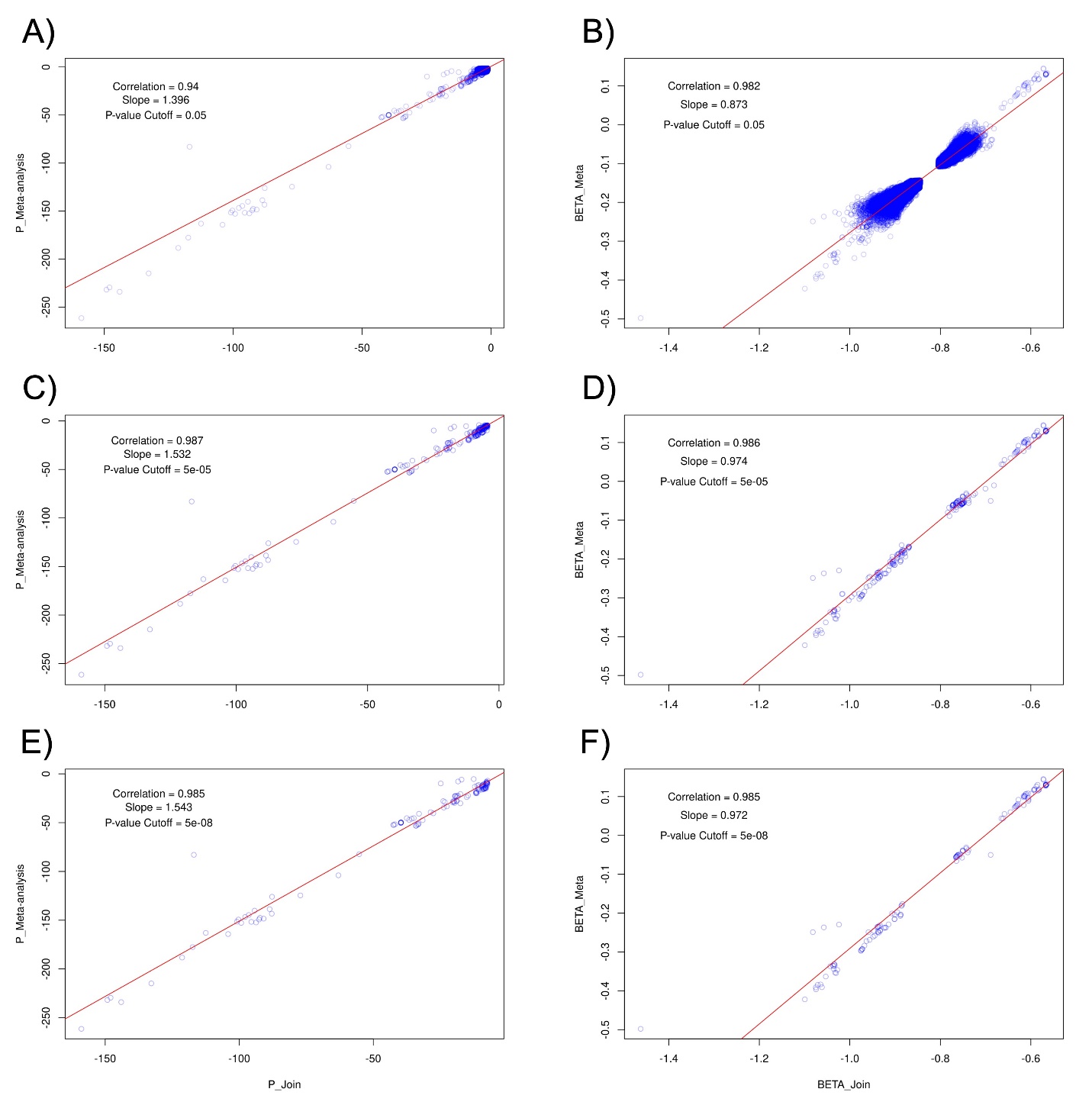

**Supplementary Figure 8: Correlation between P-values and BETA from joint-analyses using Z-scores and meta-analysis using log10 phenotype level as phenotype in 8 cohorts.** For all the cohorts having genotype and phenotype data (N=8), we performed the GWAS using log10 of raw phenotype values (e.g., PIB, Centiloid, or SUVR) for each cohort individually and then perform the SE-based meta-analysis using METAL. We also performed the joint analysis of all 8 cohorts using z-score as the quantitative phenotype. Panel A) and B) shows the correlation between the P-values and BETA, respectively, for both these analyses. Panel C) and D) also represent the same correlation plots for SNPs passing the suggestive significance threshold (P = 1 × 10−5). Panel E) and F) shows the correlation of P-value and BETA for SNPs passing the genome-wide significance thresholds (P = 5 × 10−8).

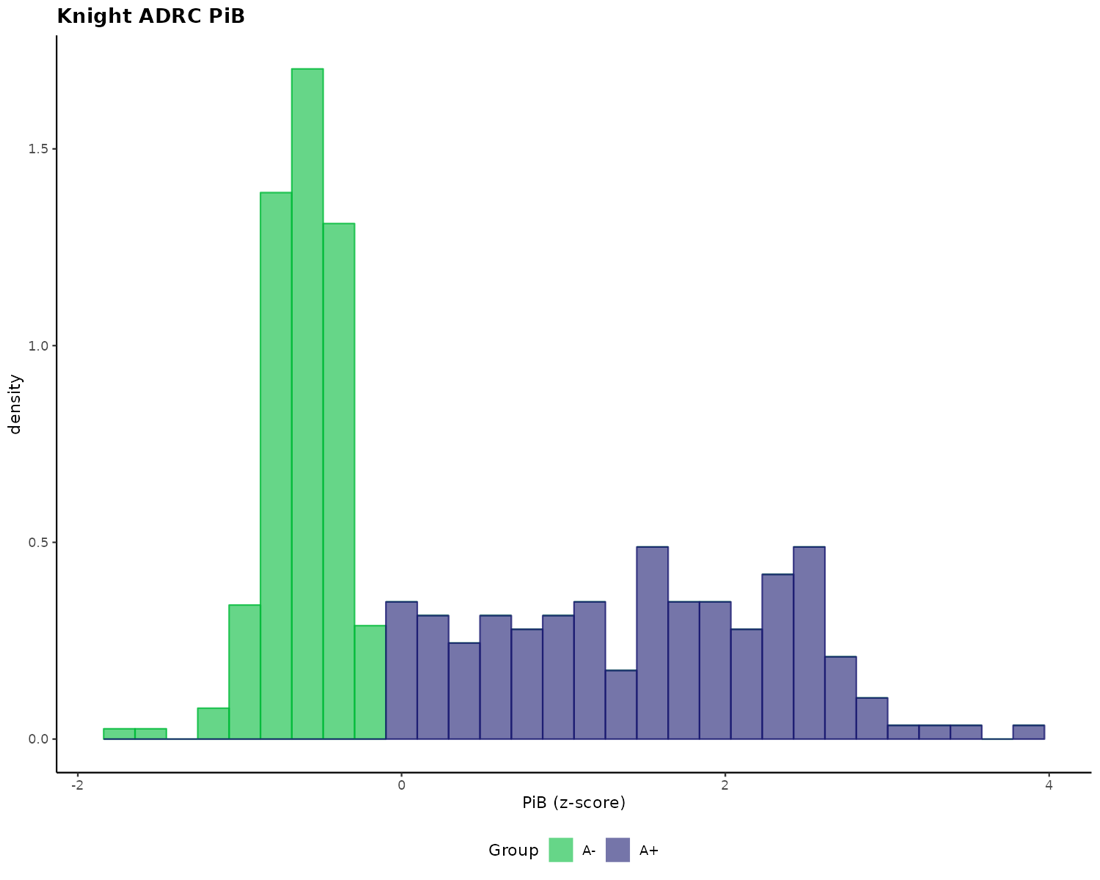

**Supplementary Figure 9: Density plot showing assumed normal distribution by z-scores based GMM approach in Knight ADRC data using PiB phenotype**. X-axis shows the z-scores values in the dataset. A+/A- denotes biomarker positivity or negativity status. Green bins represent samples assigned A- (Amyloid negative status) and bins colored purple represents samples assigned A+ (Amyloid positive status).

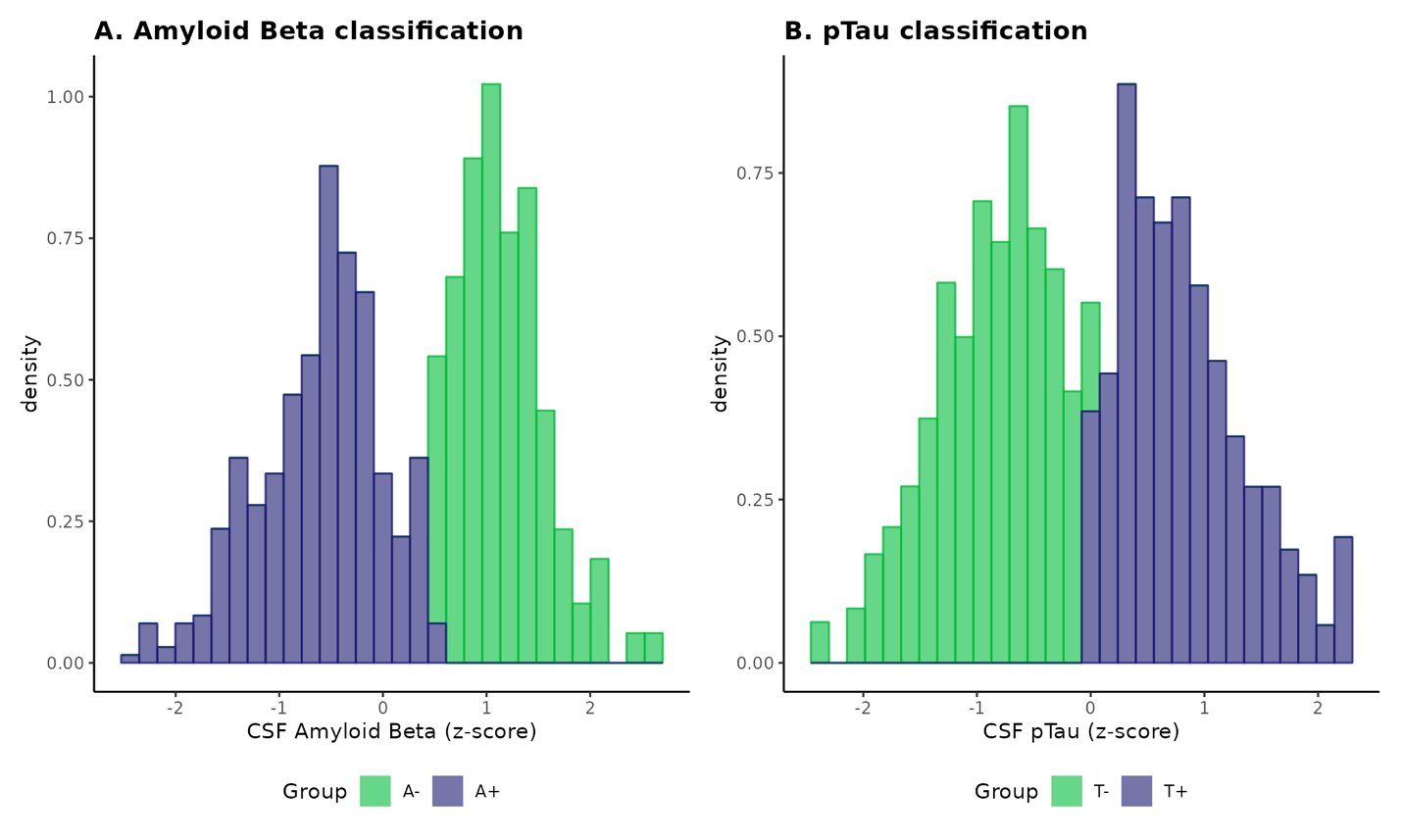

**Supplementary Figure 10: Density plot showing assumed normal distribution by z-scores based GMM approach in external validation cohort data using CSF Amyloid Beta and pTau levels.** X-axis shows the z-scores values in the dataset. A+/A- or T+/T-denotes biomarker positivity or negativity status. Green bins represent samples assigned either A- (Amyloid negative) status or T- (Tau negative) and bins colored purple represents samples assigned either A+ (Amyloid positive status) or T+ (Tau positive) status.
